# Circadian Rhythms of Perineuronal Net Composition

**DOI:** 10.1101/2020.04.21.053751

**Authors:** Harry Pantazopoulos, Barbara Gisabella, Lindsay Rexrode, David Benefield, Emrah Yildiz, Phoebe Seltzer, Jake Valeri, Gabriele Chelini, Anna Reich, Magdalena Ardelt, Sabina Berretta

## Abstract

Perineuronal Nets (PNNs) are extracellular matrix (ECM) structures that envelop neurons and regulate synaptic functions. Long thought to be stable structures, PNNs have been recently shown to respond dynamically during learning, potentially regulating the formation of new synapses. We postulated that PNNs vary during sleep, a period of active synaptic modification. Notably, PNN components are cleaved by matrix proteases such as the protease cathepsin-S. This protease is diurnally expressed in the mouse cortex, coinciding with dendritic spine density rhythms. Thus, cathepsin-S may contribute to PNN remodeling during sleep, mediating synaptic reorganization. These studies were designed to test the hypothesis that PNN numbers vary in a diurnal manner in the rodent and human brain, as well as in a circadian manner in the rodent brain, and that these rhythms are disrupted by sleep deprivation. In mice, we observed diurnal and circadian rhythms of PNNs labeled with the lectin wisteria floribunda agglutinin (WFA+PNNs) in several brain regions involved in emotional memory processing. Sleep deprivation prevented the daytime decrease of WFA+ PNNs and enhances fear memory extinction. Diurnal rhythms of cathepsin-S expression in microglia were observed in the same brain regions, opposite to PNN rhythms. Finally, incubation of mouse sections with cathepsin-S eliminated PNN labeling. In humans, WFA+PNNs showed a diurnal rhythm in the amygdala and thalamic reticular nucleus (TRN). Our results demonstrate that PNNs vary in a circadian manner and this is disrupted by sleep deprivation. We suggest that rhythmic modification of PNNs may contribute to memory consolidation during sleep.

## Introduction

Perineuronal Nets (PNNs) are extracellular matrix (ECM) structures surrounding subpopulations of neurons. PNNs form during the end of critical periods of plasticity, marking their closure by conferring an adult form of restricted plasticity (Pizzorusso et al., 2002; Gogolla et al., 2009; Mauney et al., 2013). Although PNNs have been historically considered stable structures, recent studies suggest they are modified during learning to allow for formation of synapses (Nagy et al., 2007; Brown et al., 2009; Ganguly et al., 2013; Banerjee et al., 2017; Slaker et al., 2018). An important line of evidence comes from studies on matrix metalloproteases, which cleave ECM components including chondroitin sulfate proteoglycans (CSPGs), key components of PNNs, and contribute to regulation of synaptic plasticity (Muir et al., 2002; Porter et al., 2005; Bajor and Kaczmarek, 2013). Expression of these proteases contributes to fear learning and memory consolidation (Brown et al., 2009; Ganguly et al., 2013). These effects are mediated through dendritic spine remodeling (Szklarczyk et al., 2002) and regulation of long-term plasticity (LTP) (Nagy et al., 2006). In addition, increasing number of studies directly show that CSPGs, and in turn PNNs, are critically involved in the regulation of synaptic plasticity (Bukalo et al., 2001). Notably, the strength of LTP has been shown to vary in a circadian manner in the hippocampus (Chaudhury et al., 2005), a region where PNN functions in regulating synaptic strength and stability have been particularly well characterized (Bukalo et al., 2001; Brakebusch et al., 2002; Geissler et al., 2013). Together, these observations support the hypothesis that PNN composition may be regulated in a circadian manner to allow for circadian rhythms in synaptic plasticity.

PNN composition is regulated by several cell types including astrocytes and neurons, which produce several of the core PNN components, along with a broad range of endogenous proteases that cleave PNN components produced primarily by astrocytes and microglia (Pantazopoulos and Berretta, 2016; Miyata and Kitagawa, 2017; Bozzelli et al., 2018). We focus on the microglial protease cathepsin-S as a first step towards identifying molecules that may contribute to circadian rhythms in PNN composition. Cathepsin-S has been shown to regulate synaptic plasticity, cleave several ECM components, and is expressed diurnally in the rodent cortex (Petanceska et al., 1996; Hayashi et al., 2013b). Furthermore, several lines of evidence suggest that microglia contribute to diurnal regulation of PNNs. Compelling data point to the role of microglia in the regulation of synaptic plasticity (Wake et al., 2013; Stevens and Schafer, 2018). Furthermore, microglial dysfunction in the hippocampus results in reduction of dendritic spines along with increased ECM expression (Bolós et al., 2018), suggesting microglia participate in degrading the ECM to allow for increased synaptic plasticity. A recent study demonstrated that pharmacological depletion of microglia prevented PNN decreases that normally occur in a mouse model of Huntington’s disease and improved memory function (Crapser et al., 2020), suggesting that microglia play a critical role in regulating PNN composition. Taken together, the current evidence suggests that cathepsin-S from microglia is an optimal candidate for contributing to modification of PNN composition to allow dynamic regulation of synaptic plasticity during sleep.

PNNs are well represented in neural circuits involved in emotion processing and critically involved in the regulation of fear and reward memories (Gogolla et al., 2009; Slaker et al., 2015; Banerjee et al., 2017; Lasek et al., 2018). Consistent with these observations, PNNs have been implicated in several brain disorders involving these regions, including schizophrenia, bipolar disorder, Alzheimer’s disease and addiction (Baig et al., 2005; Morawski et al., 2010; Pantazopoulos et al., 2010a; Mauney et al., 2013; Xue et al., 2014; Pantazopoulos et al., 2015; Slaker et al., 2015; Steullet et al., 2017; Blacktop and Sorg, 2018). Several of these disorders also show altered sleep and circadian rhythms (Lim et al., 2013; McClung, 2013; Wang et al., 2015; Manoach et al., 2016; Pantazopoulos et al., 2017). Thus, diurnal modulation of PNNs has a broad range of implications for psychiatric disorders and memory processing.

We tested the hypothesis that PNNs vary in a circadian manner, and that these rhythms are disrupted by sleep deprivation. In mice, we first assessed densities of PNNs across the 24-hour cycle in brain regions involved in emotional memory processing and implicated in psychiatric disorders (Vyas et al., 2002; Sartorius et al., 2010; Li et al., 2011; Mahan and Ressler, 2012; Mauney et al., 2013; Meyer et al., 2014; Pantazopoulos et al., 2017; Wells et al., 2017). We then assessed the relationship between PNNs and sleep by testing the effect of sleep deprivation on PNN densities in several regions including the hippocampus, a brain region in which diurnal differences in LTP were reported (Chaudhury et al., 2005).

As a first step in testing the hypothesis that matrix proteases are involved in regulating PNN rhythmicity, we characterized rhythms of cathepsin-S expression. We then demonstrated that cathepsin-S impacts PNN integrity by incubating mouse sections in active cathepsin-S enzyme. Finally, we tested the hypothesis that PNN rhythmicity is conserved in humans, focusing on the amygdala and thalamic reticular nucleus (TRN), two regions in which PNN deficits in schizophrenia and bipolar disorder were reported (Pantazopoulos et al., 2010b; Pantazopoulos et al., 2015; Steullet et al., 2017).

## Methods and Materials

### Antibodies and Lectin Labeling

#### Wisteria Floribunda Agglutinin (WFA)

WFA, (catalogue #B-1355, Vector Labs, Burlingame, CA), a lectin isolated from seeds of *wisteria floribunda*, binds specifically to N-acetyl-D-galactosamine on the terminal end of chondroitin sulfate (CS) chains, with a preference for beta glycosidic linkage (Kurokawa et al., 1976). The specificity of WFA as a marker for these macromolecules is supported by extensive literature, including ablation of labeling following CS enzymatic digestion (Galtrey and Fawcett, 2007; Pantazopoulos et al., 2010a).

#### Cathepsin-S (E-3)

Cathepsin-S E-3 (sc-271619, Santa Cruz Biotechnology Inc., Dallas, TX) is a mouse monoclonal antibody raised against a peptide matching amino acids 302-331 at the C-terminus of human cathepsin-S, shown to detect the 24 kDa form of cathepsin-S (sc-271619 data sheet, Santa Cruz Biotechnology Inc., Dallas, TX).

#### IBA1

IBA1 (019-19741, FUJIFILM Wako Chemicals USA, Richmond, VA) is a rabbit polyclonal antibody raised against a synthetic peptide to the C-terminus of IBA1, shown to detect the 17 kDa form of IBA1 in rat and mouse brain samples (019-19741data sheet, FUJIFILM Wako Chemicals USA, Richmond, VA).

### Immunocytochemistry (Mouse samples)

Free-floating tissue sections were carried through antigen retrieval in citric acid buffer (0.1 M citric acid, 0.2 M Na2HPO4) heated to 80 degrees °C for 30 minutes, and incubated in biotinylated WFA lectin (cat#B-1355, Vector Labs) or the mouse monoclonal primary antibody anti-cathepsin-S (1:500, sc-271619, Santa Cruz Biotechnology Inc.) for 48 hours, and subsequently in biotinylated secondary antibody (horse anti-goat IgG; 1:500; Vector Labs, Inc. Burlingame, CA), followed by streptavidin conjugated with horse-radish peroxidase for two hours (1:5000 μl, Zymed, San Francisco, CA), and, finally, in nickel-enhanced diaminobenzidine/ peroxidase reaction (0.02% diaminobenzidine, Sigma-Aldrich, 0.08% nickel-sulphate, 0.006% hydrogen peroxide in PB). All solutions were made in PBS with 0.2% Triton X (PBS-Tx) unless otherwise specified. Immunostained sections were mounted on gelatin-coated glass slides, coverslipped and coded for blinded quantitative analysis. All sections included in the study were processed simultaneously within the same session to avoid procedural differences. Omission of the primary or secondary antibodies did not result in detectable signal, and pre-absorption of mouse anti cathepsin-S with 300 nanograms of active human cathepsin-S (SRP0292, Sigma-Aldrich, St. Louis, MO) did not result in detectable immunolabeling signal.

### Dual Antigen Immunofluorescence

Sections were co-incubated in primary antibodies (cat-S, l:500 µl, rabbit anti-IBA1, l:1000 µl (FUJIFILM Wako, cat#019-19741); in 2% bovine serum albumin (BSA) for 72 hr at 4 °C. This step was followed by 4 hour incubation at room temperature in Alexa Fluor goat anti-mouse 647 (1:300 µl; A-21235, Invitrogen, Grand Island, NY) and donkey anti-rabbit 555 (1:300 µl; A-32794, Invitrogen, Grand Island, NY), 10 minutes incubation in DAPI 1:16000 in 0.1M PB, followed by 10 minutes in 1 mM CuSO4 solution (pH 5.0) to block endogenous lipofuscin autofluorescence (Schnell et al., 1999). Sections were mounted and coverslipped using Dako mounting media (S3023, Dako, North America, Carpinteria, CA).

### Immunocytochemistry (Human Samples)

Free-floating tissue sections were carried through antigen retrieval in citric acid buffer (0.1 M citric acid, 0.2 M Na2HPO4) heated to 80 degrees °C for 30 minutes, and incubated in biotinylated WFA lectin (cat#B-1355, Vector Labs) for 48 hours, followed by streptavidin conjugated with horse-radish peroxidase for two hours (1:5000 μl, Zymed, San Francisco, CA), and, finally, in nickel-enhanced diaminobenzidine/ peroxidase reaction (0.02% diaminobenzidine, Sigma-Aldrich, 0.08% nickel-sulphate, 0.006% hydrogen peroxide in PB). All solutions were made in PBS with 0.5% Triton X (PBS-Tx) unless otherwise specified. Immunostained sections were mounted on gelatin-coated glass slides, coverslipped and coded for blinded quantitative analysis. All sections included in the study were processed simultaneously within the same session to avoid procedural differences. Omission of the WFA lectin or HRP conjugated streptavidin did not result in detectable signal.

### Data Collection (Mouse)

In mouse brain samples, serial sections containing the hippocampus, infralimbic cortex, prelimbic cortex, TRN, and habenula were quantified using a Leica microscope interfaced with Bioquant Nova Prime v6.0, (R&M Biometrics, Nashville, Tennessee). Borders of each region were defined according to the Allen Brain Atlas and traced under 4x magnification. Each traced region was systematically scanned through the full x, y, and z-axes under 40x magnification to count each WFA+ PNN or cathepsin-S immunoreactive (IR) microglial cell.

Dual immunofluorescence sections labeled for cathepsin-S and IBA1 from three adult male mice housed in a standard light-dark cycle (4 sections per mouse) and sacrificed at ZT6 were quantified using Stereo-Investigator Image Analysis System (v.10.0; MBF Biosciences, Williston, VT), interfaced with an Olympus BX-61 microscope. Cathepsin-S immunoreactive cells were distinguished from cathepsin-S immunoreactive blood vessels by the presence or absence of DAPI stained nuclei.

### Data Collection (Human)

In human postmortem samples, total numbers and numerical densities of PNNs labeled with WFA were quantified using stereology based sampling (Pantazopoulos et al., 2007; Dorph-Petersen and Lewis, 2011) in the amygdala and TRN in a cohort of postmortem brain samples from human subjects (14 amygdala, 15 TRN subjects). WFA labeled (WFA+) PNNs were counted in the lateral (LN), basal (BN), accessory basal (AB) and cortical (CO) nuclei of the amygdala, and in TRN using a Zeiss Axioskop-2 Plus interfaced with Stereo-Investigator 6.0 (Microbrightfield Inc., Willinston, VT). Intra-rater (H.P. and M.A.) reliability of at least 95% was established before formal data collection and reassessed regularly. The borders of amygdala nuclei were traced and confirmed in adjacent Nissl stained sections according to cytoarchitectonic criteria described by Amaral et al, 1992 and Sims and Williams, 1990 (Sims and Williams, 1990; Amaral et al., 1992). The nomenclature adopted was used by Sorvari et 1995 (Sorvari et al., 1995). The central, medial and anterior nuclei could not be quantified because their dorso-medial portion was damaged in several samples. The borders of the TRN were identified according to specific landmarks, such as the internal capsule laterally and the subthalamic nucleus ventromedially. Each traced region was systematically scanned through the full x, y, and z-axes to count each WFA labeled PNN over complete sets of serial sections (6-10 sections) representing the whole extent of the amygdala from each subject (section interval 1040 µm). Outcome measures were plotted by time of death for each subject to analyze potential diurnal fluctuations using approaches reported previously in postmortem studies (Monk et al., 1997; Dumont et al., 1999; Zhou et al., 2001; Hofman, 2003; Iwata et al., 2013; Li et al., 2013; Schmal et al., 2013; Pantazopoulos et al., 2016).

### Statistical Analysis

Differences between groups relative to the main outcome measures were assessed for statistical significance using stepwise linear regression (ANCOVA). Logarithmic transformation was uniformly applied to all human data values because data were not normally distributed. Statistical analyses were performed using JMP PRO v14 (SAS Institute Inc., Cary, NC). Average daily wheel-running activity was included as a covariate for all mouse studies. Time of death (TOD) was obtained from the death certificate for each subject and tested for potential effects on outcome measures. TOD was also used to divide subjects into subjective day (s-Day TOD, 06:00-17:59 hours) and subjective night (s-Night, 18:00-05.59 hours) groups on the basis of previous literature indicating diurnal fluctuations in the amygdala of humans and mice (Berelowitz et al., 1981; Arnold et al., 1982; Rubinow, 1986). Effects of TOD on outcome measures were analyzed using two steps: 1) Subjects were divided into s-Day vs. s-Night groups for comparisons using stepwise linear regression analysis 2) We used quartic regression analysis on plots of **Nt** of WFA labeled PNNs by TOD for each group according to methods used to detect similar relationships in postmortem studies (Zhou et al., 2001; Hofman, 2003; Li et al., 2013). Quartic regression models were used as described previously (Pantazopoulos et al., 2017) to fit expression patterns reported in the mouse and human amygdala consisting of two peaks and two troughs (Albrecht et al., 2013; Pantazopoulos et al., 2017).

### Numerical Densities (Mouse Samples)

Numerical densities were calculated as **Nd= ∑N / ∑V** where **N =** sum of all PNNs counted in each region for each animal, and **V** is the volume of each region per animal, calculated as **V= ∑a • z**, where z is the thickness of each section (30 μm) and a is area in μm2. Rhythmic relationships of PNNs and cathepsin-S in mice were analyzed by plotting means and standard deviation per each time point across the 24-hour cycle, as conducted in previous studies (Lamont et al., 2005; Segall et al., 2009; Harbour et al., 2014).

### Numerical Densities and Total Numbers Estimates (Human Samples)

Total number (**Nt**) of WFA labeled PNNs was calculated as **Nt = i • ∑n** where ∑**n** = sum of the cells counted in each subject, and **i** is the section interval (i.e. number of serial sections between each section and the next within each compartment=26) as described previously (Berretta et al., 2007). Numerical densities were calculated as **Nd= ∑N / ∑V** where **V** is the volume of each amygdala nucleus or the TRN, calculated as **V= z • ssf • ∑ a** where ***z*** is the thickness of the section (40 µm), ***ssf*** is the section sampling fraction (1/26; i.e. number of serial sections between each section within a compartment) and **a** is the area of the region of interest.

### Animals

Adult male wild type C57/BL6 mice housed in individual wheel-running cages in a 12:12 light:dark (LD) cycle were used to examine diurnal rhythms of PNN composition. Three male C57/BL6 mice were sacrificed every 4 hours across the 24-hour cycle at zeitgeber time (ZT) 0, 4, 8, 12, 16, and 20. A separate set of adult male C57/BL6 mice were used to test for circadian rhythms of PNN composition. Mice were housed in a 12:12 LD cycles for 4 weeks, followed by three full 24-hour cycles in constant darkness, then sacrificed every 4 hours at circadian time (CT) 0, 4, 8, 12, 16, and 20), 3 mice per timepoint. Wheel-running actigraphs were used to determine individual CT times for sacrificing animals housed in constant darkness. Activity onset over three 24-hour cycles was used to predict CT time in the 4th cycle during which animals were sacrificed. All animals in the constant darkness study were sacrificed under dim red light conditions. Circadian rhythm of each mouse was monitored with ClockLab (Actimetrics, Wilmette, IL) using wheel-running activity data. Mice were sacrificed using cervical dislocation in the light or in the dark using a dim red light, depending on lighting conditions at time of sacrifice. Mice were perfused intracardially with 4% PFA and brains were stored in 0.1 M PB with 0.1% Na Azide and 30% sucrose. Brains were then sliced into serial 30 μm brain sections on an American Optical freezing microtome. The housing and treatment of experimental animals were approved by the University of Mississippi Medical Center Institutional Animal Care and Use Committee and followed guidelines set by the National Institutes of Health.

### Human Subjects and Tissue Processing

Tissue blocks containing the whole amygdala or thalamus from 15 donors were obtained from the Harvard Brain Tissue Resource Center, McLean Hospital, Belmont, MA (*Table 1*). Diagnoses were made by two psychiatrists on the basis of retrospective review of medical records and extensive questionnaires concerning social and medical history provided by family members. A neuropathologist examined several regions from each brain for a neuropathology report. The cohort for this study did not include subjects with evidence for gross and/or macroscopic brain changes, or clinical history consistent with cerebrovascular accident or other neurological disorders. Subjects with Braak & Braak stages III or higher were not included. Subjects had no significant history of psychiatric illness, or substance dependence, other than nicotine and alcohol, within 10 years from death.

**Table 1.** Thalamic Reticular Nucleus Sample Demographic and Descriptive Characteristics.

| Case | Age | Sex | Cause of death | brain weight (g) | PMI (hrs) | Hemisphere | TOD |
| --- | --- | --- | --- | --- | --- | --- | --- |
| S05735 | 74 | F | Cancer (C) | 1145 | 12.2 | L | 14.00 |
| S16022 | 68 | F | Cardiac Arrest (A) | 1330 | 14.75 | R | 09.30 |
| S13845 | 37 | M | Electrocution (A) | 1460 | 18.75 | R | 21.00 |
| S14247 | 72 | M | Cardiac Arrest (A) | 1560 | 28.2 | R | 07.35 |
| S10160 | 85 | M | Cancer (C) | 1225 | 20.3 | L | 05.30 |
| S18228 | 78 | F | Cancer (C) | 1100 | 23.9 | L | 05.00 |
| S07594 | 95 | F | Unknown | 1350 | 7.1 | R | 14.50 |
| S06087 | 69 | F | Unknown | 1280 | 25.2 | R | 10.16 |
| S07749 | 61 | M | Unknown | 1280 | 10.1 | R | 12.30 |
| S07429 | 68 | F | Unknown | 1230 | 24.8 | R | 19.45 |
| S14342 | 70 | F | Cardiac Arrest (A) | 1245 | 18.0 | R | 07.29 |
| S08987 | 53 | F | Cancer (C) | 1330 | 24.0 | R | 08.32 |
| S11774 | 74 | M | Cardiac Arrest (A) | 1490 | 15.81 | R | 08.41 |
| S03774 | 70 | M | Aortic Aneurysm (A) | 1400 | 17.3 | R | 20.46 |
| S17165 | 58 | F | COPD (C) | 1345 | 17.8 | R | 00.35 |
| Total/mean ± SD | 68.8±13.5 | 9F 6M |  | 1318.0±125.1 | 18.5±6.0 | 3L 12R |  |

**Table 2.** Amygdala Sample Demographic and Descriptive Characteristics.

| Case | Age | Sex | Cause of death | brain weight (g) | PMI (hrs) | Hemisphere | TOD |
| --- | --- | --- | --- | --- | --- | --- | --- |
| S90122 | 70 | M | Cardiac Arrest (A) | 1360 | 23.2 | L | 12.17 |
| S23073 | 52 | M | Cardiac Arrest (A) | - | 32.1 | L | 03.07 |
| S12827 | 71 | M | Cardiac Arrest (A) | 1580 | 24.0 | L | 10.10 |
| S13845 | 37 | M | Electrocution (A) | 1460 | 18.75 | R | 21.00 |
| S07340 | 65 | M | Cardiac Arrest (A) | 1240 | 17.3 | L | 06.45 |
| S08987 | 53 | F | Cancer (C) | 1330 | 24.0 | R | 08.32 |
| S30877 | 62 | M | Cardiac Arrest (A) | 1300 | 29.2 | L | 21.18 |
| S03774 | 70 | M | Aortic Aneurysm (A) | 1400 | 17.3 | R | 20.46 |
| S17232 | 58 | M | COPD (C) | 1066 | 19.3 | R | 16.08 |
| S14247 | 72 | M | Cardiac Arrest (A) | 1560 | 28.2 | R | 07.35 |
| S05735 | 74 | F | Cancer (C) | 1145 | 12.2 | L | 14.00 |
| S16022 | 68 | F | Cardiac Arrest (A) | 1330 | 14.75 | R | 09.30 |
| S10160 | 85 | M | Cancer (C) | 1225 | 20.3 | L | 05.30 |
| S18228 | 78 | F | Cancer (C) | 1100 | 23.9 | L | 05.00 |
| Total/mean ± SD | 65.4±12.2 | 5F 9M | | 1315.0±161.3 | 21.8±5.7 | 8L 6R | |

Abbreviations: A, acute death, no prolonged agonal period; C, chronic, prolonged agonal period; COPD, Chronic Obstructive Pulmonary Disease; PMI, postmortem time interval

Tissue blocks were dissected from fresh brains and post-fixed in 0.1M PB containing 4% paraformaldehyde and 0.1M Na azide at 4°C for 3 weeks, cryoprotected at 4°C for 3 weeks (30% glycerol, 30% ethylene glycol and 0.1% Na azide in 0.1M PB), embedded in agar, and pre-sliced in 2 mm coronal slabs using an Antithetic Tissue Slicer (Stereological Research Lab., Aarhus, Denmark). Each slab was exhaustively sectioned using a freezing microtome (American Optical 860, Buffalo, NY). Sections were stored in cryoprotectant at −20°C. Using systematic random sampling criteria, sections through the amygdala were serially distributed in 26 compartments (40 µm thick sections; six-ten sections/compartment; 1.04 mm section separation within each compartment). All sections within one compartment/subject were selected for histochemistry (i.e., WFA), thus respecting the ‘equal opportunity’ rule (Coggeshall and Lekan, 1996; Gundersen et al., 1999).

### Sleep Deprivation

Adult male wild type C57/Bl6 mice housed in 12:12 LD cycle were used for sleep deprivation experiments. Mice were either sleep deprived using gentle handling for 5 hours from lights on (7 AM) to 12 PM (n=12) or handled during the dark phase for 5 hours the night before (controls; n=12), to control for potential confounding effects of handling on the outcome measures. Mice were sacrificed immediately following 5 hours of sleep deprivation, and control mice were sacrificed at the same time (ZT 5: 12 PM). Mice were perfused intracardially with 4% PFA and brains were stored in 0.1 M PB with 0.1% Na Azide and 30% sucrose. Brains were then sliced into serial 30 μm brain sections on an American Optical freezing microtome. WFA labeling was used to quantify PNNs in the habenula, prefrontal cortex, amygdala, thalamus, and hippocampus using stereology-based sampling methods.

### Mouse Auditory Fear Conditioning

Auditory contextual fear conditioning was conducted as described previously (Gisabella et al., 2016). Mice were placed in a fear conditioning box at ZT0 (7am), (64cm wide ∼ 73 cm deep ∼ 68cm high) (Med Associates, St. Albans, VT, USA) placed in a larger, sound-attenuating chamber. Precisely four mice will be placed in 4 boxes chamber (one mouse for each box) at the same time for experimental comparison. Mice remained in the chamber for three minutes before delivery of four tones, each of 10 seconds duration (85dB, 10kHz). Each tone was followed by a footshock lasting 2 seconds (0.8 mA amplitude) pairings were administered (60–200 s variable ITI) (total time mice spent inside the chamber was 15-18mins). After the test the mice were placed back into normal housing (4 control mice, or sleep deprived for 5 hours before being placed back into normal housing (4 mice).

Mice were returned to the context on the second day for an extinction session (10 min total inside the chamber with no shock and tone) then placed back in their cages. Mice were placed in a novel context on the third day at 7am for auditory fear extinction inside the chamber for a total time of 5 minutes (3 min pre; 60 s, 85dB, 10kHz tone; 60 s post-tone period). Low freezing prior to the onset of tone presentation indicated that animals did not generalize fear to the novel context, and also enabled us to conclude that freezing observed during the tone was evoked specifically by the tone. Freezing behavior was defined as periods of at least 1 s with the complete absence of movement except breathing; it was measured with manual scoring. The percent of time spent freezing during intervals of interest was quantified, and these results were analyzed using analysis of variance (ANOVA). Post hoc Fisher’s PLSD tests were performed after a significant omnibus F-ratio.

### Cathepsin-S PNN Digestion

Free floating mouse brain sections were incubated with 300 nanograms of active human cathepsin-S (SRP0292, Sigma-Aldrich, St. Louis, MO), in activation buffer containing 1.8 mM DTT, 1.8mM EDTA, 1% BSA, 12mM citric acid, 43 mM Na2HPO4 at 37 ºC for either 3 hours or 24 hours. Control sections were incubated in activation buffer (1.8 mM DTT, 1.8mM EDTA, 1% BSA, 12mM citric acid, 43 mM Na2HPO4) at 37 ºC in parallel. Following cathepsin-S incubation, sections were labeled with WFA and WFA+ PNNs were quantified in the hippocampus as described above.

## Results

We use the chronobiology term ‘circadian’ to refer to rhythms observed in constant darkness, regulated by endogenous circadian processes in the absence of environmental signals that can entrain rhythms such as light-dark cycles. We use the term ‘diurnal’ to refer to rhythms observed in light-dark cycles, which may reflect immediate responses to environmental cycles rather than true circadian rhythms.

### Diurnal Rhythms of PNNs in the Mouse Brain

In a cohort of adult male C57/BL6 mice, housed in a 12:12 LD cycle, we observed diurnal rhythms in the density of WFA+ PNNs in the hippocampus, amygdala, prefrontal cortex, habenula, and TRN (Figs. 1-5). WFA+ PNN rhythms in the hippocampal sectors displayed consistent peaks at ZT20, and troughs at ZT6 across hippocampal sectors (Fig. 1). WFA+ PNN density rhythms in the amygdala were similar to hippocampal rhythms, with peaks at ZT20 and troughs at ZT8 across amygdala nuclei (Fig.2). Similar relationships were observed in the prefrontal cortex, with WFA+ PNN densities displaying peaks at approximately ZT0 and troughs at ZT8 (Fig.3). WFA+ PNN density rhythms in the habenula (Fig.4) and TRN (Fig. 5) also displayed consistent diurnal rhythms, with peaks at approximately ZT20 and troughs at ZT8. ANCOVA models testing the main effect of ZT time and the effect of average daily amount of wheel running activity showed significant effects of ZT time on WFA+PNN densities in all regions examined (Table 3). In comparison, average daily amount of wheel-running activity showed a significant effect on densities of WFA+PNNs in only the central amygdala and thalamic reticular nucleus (Table 3).

**Table 3.** Summary Table of ZT time and Average Daily Running Activity Effects on WFA+ PNN and CatS-immunoreactive Cell Densities.

|  | ZT time<br>F-ratio | ZT time<br>p-value | Avg daily Wheel<br>running activity<br>F-ratio | Avg daily wheel<br>running activity<br>p-value |
| --- | --- | --- | --- | --- |
| CA1 PNNs | **12.51** | **0.004** | 0.01 | 0.92 |
| CA2/3 PNNs | **20.21** | **<0.0001** | 4.07 | 0.07 |
| CA4 PNNs | **26.55** | **<0.0001** | 23.71 | 0.12 |
| DG PNNs | **7.38** | **0.003** | 0.02 | 0.91 |
| Lateral amygdala PNNs | **9.16** | **0.001** | 0.02 | 0.98 |
| Basolateral amygdala PNNs | **10.81** | **0.0007** | 1.72 | 0.22 |
| Central amygdala PNNs | **48.66** | **<0.0001** | **18.15** | **0.002** |
| TRN PNNs | **4.69** | **0.02** | **5.71** | **0.03** |
| Lateral habenula PNNs | **10.25** | **0.0009** | 0.88 | 0.37 |
| Medial Habenula PNNs | **5.31** | **0.01** | 0.07 | 0.78 |
| IL Superficial PNNs | **14.72** | **0.001** | 0.03 | 0.96 |
| IL Deep PNNs | **10.94** | **0.003** | 0.88 | 0.38 |
| PL Superficial PNNs | **28.16** | **0.0002** | 2.51 | 0.16 |
| PL Deep PNNs | **31.77** | **<0.001** | 0.94 | 0.37 |
| CA1 CatS | **9.48** | **0.008** | 2.29 | 0.18 |
| CA2/3 CatS | **5.88** | **0.02** | 1.12 | 0.33 |
| CA4 CatS | **6.79** | **0.01** | 3.01 | 0.13 |
| DG CatS | **54.10** | **<0.0001** | 5.59 | 0.06 |
| Lateral amygdala CatS | **6.01** | **0.02** | 0.22 | 0.65 |
| Basolateral amygdala CatS | **26.25** | **0.0002** | 3.95 | 0.09 |
| Central amygdala CatS | **14.21** | **0.001** | 0.02 | 0.88 |
| IL Superficial CatS | **8.20** | **0.05** | 0.17 | 0.71 |
| IL Deep CatS | **7.67** | **0.04** | 0.50 | 0.52 |
| PL Superficial CatS | **28.21** | **0.009** | 0.18 | 0.70 |
| PL Deep CatS | **93.77** | **0.002** | 0.009 | 0.92 |

Values represent F-ratios and p values derived from ANCOVA models testing effects of ZT time and average daily wheel-running activity on WFA+ PNN densities and cathepsin-S immunoreactive cell densities. Statistically significant differences are indicated in BOLD = $p < 0.05$ .

**Figure 1:**
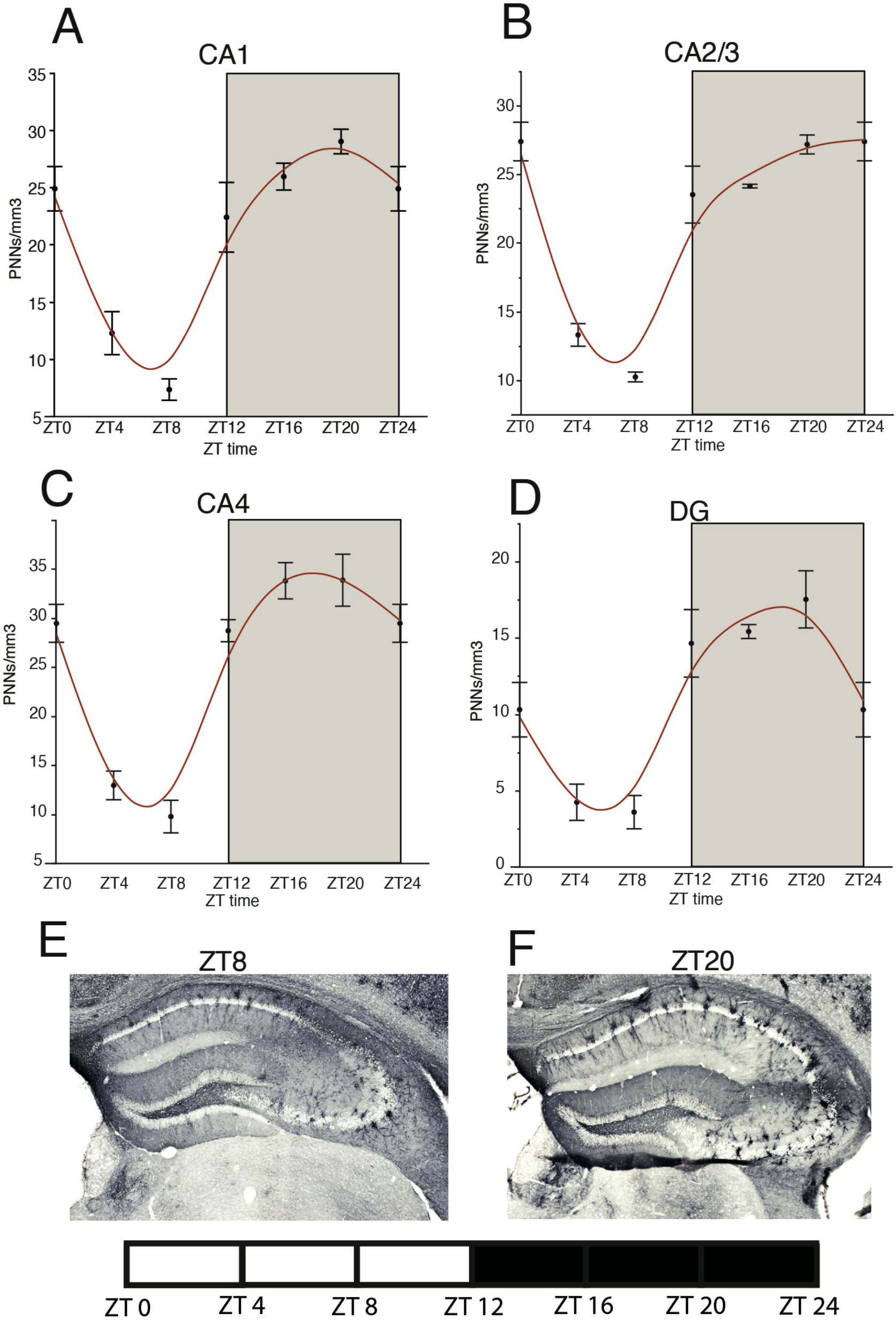
Diurnal Rhythms of Perineuronal Nets in the Mouse Hippocampus. Analysis of WFA+ PNNs across the 24 hour cycle in male mice housed in a 12:12 light/dark cycle revealed a diurnal rhythm of WFA+ PNNs in hippocampal sectors CA1 (A) CA2/3 (B), CA4 (C) and the DG (D) with peaks at ∼ZT20 and troughs at ∼ZT8. Error bars represent standard deviation. Representative low magnification images of WFA labeling in the mouse hippocampus at ZT 8 (E) and ZT 20 (F).

**Figure 2:**
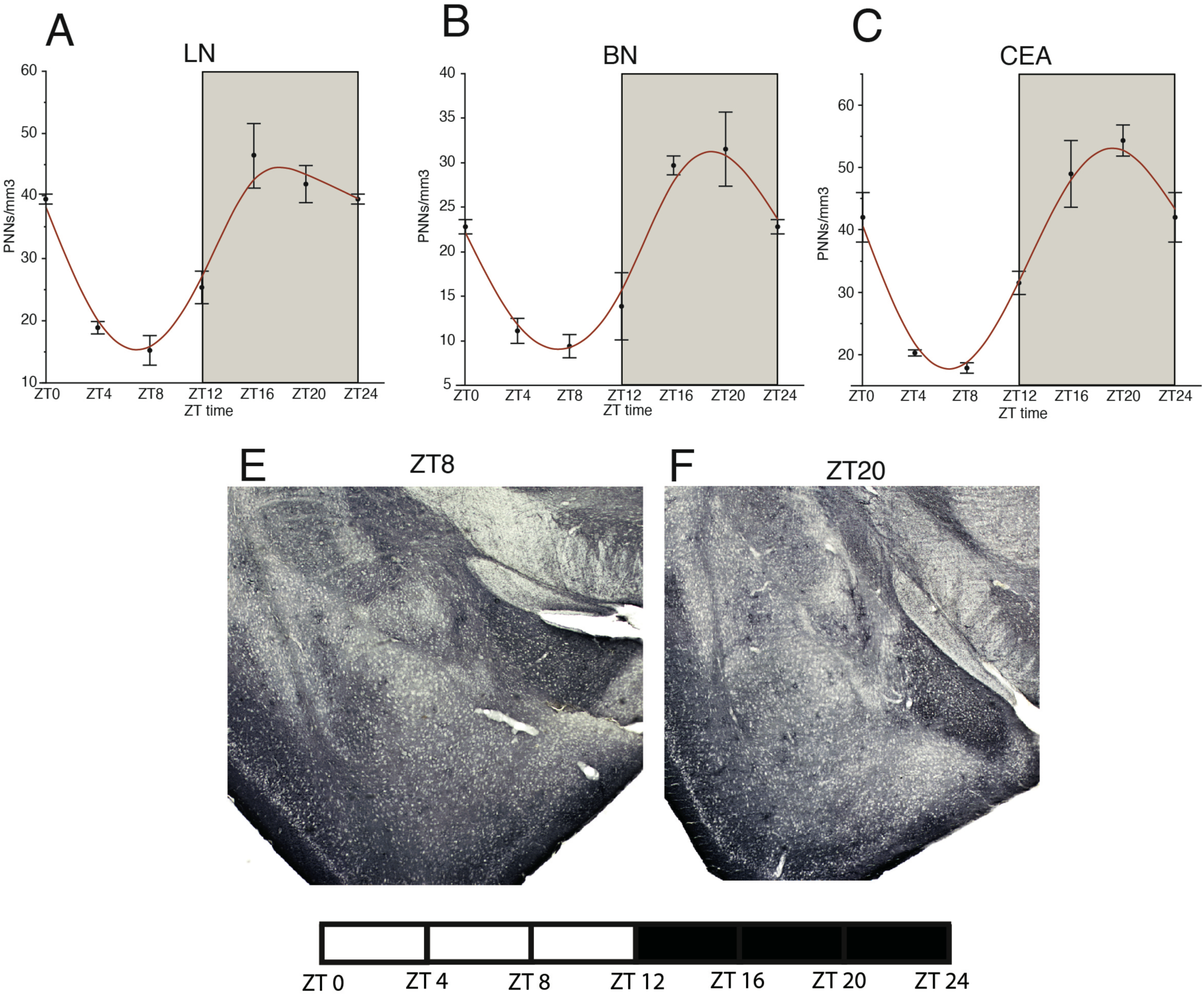
Diurnal Rhythms of Perineuronal Nets in the Mouse Amygdala. Diurnal rhythms of WFA+ PNNs were observed in the lateral amygdala (A) basal amygdala (B), and central amygdala (C) with peaks at ∼ZT20 and troughs at ∼ZT8. Error bars represent standard deviation. Representative low magnification images of WFA labeling in the mouse amygdala at ZT 8 (D) and ZT 20 (E).

**Figure 3:**
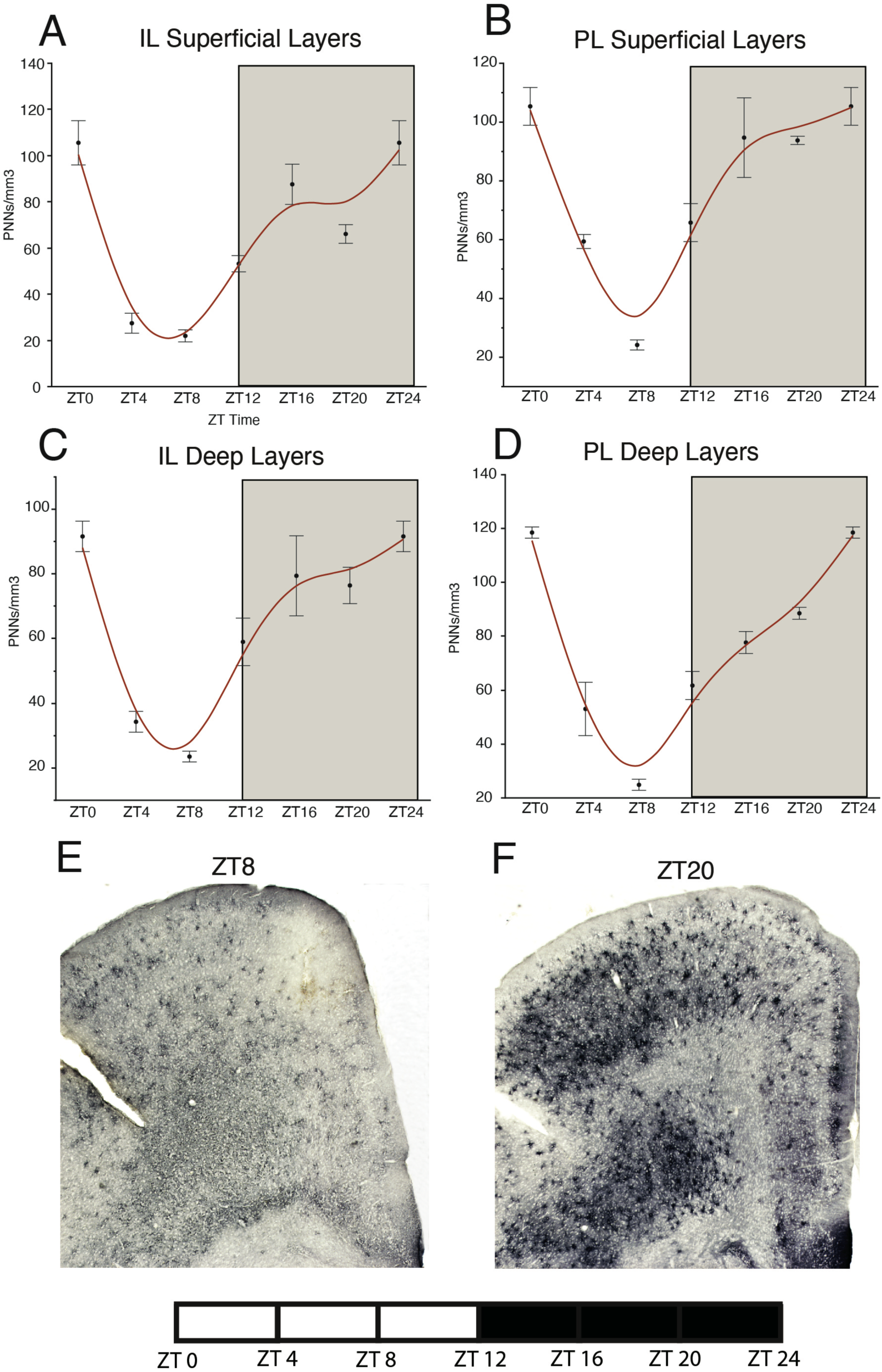
Diurnal Rhythms of Perineuronal Nets in the Mouse Prefrontal Cortex. Diurnal rhythms of WFA+ PNNs were observed in the infralimbic superficial (A) prelimbic superficial (B), infralimbic deep (C) and prelimbic deep (D) layers of the mouse, with peaks at ∼ZT 0 and troughs at ∼ZT 8. Error bars represent standard deviation. Representative low magnification images of WFA labeling in the mouse prefrontal cortex at ZT 8 (E) and ZT 20 (F).

**Figure 4:**
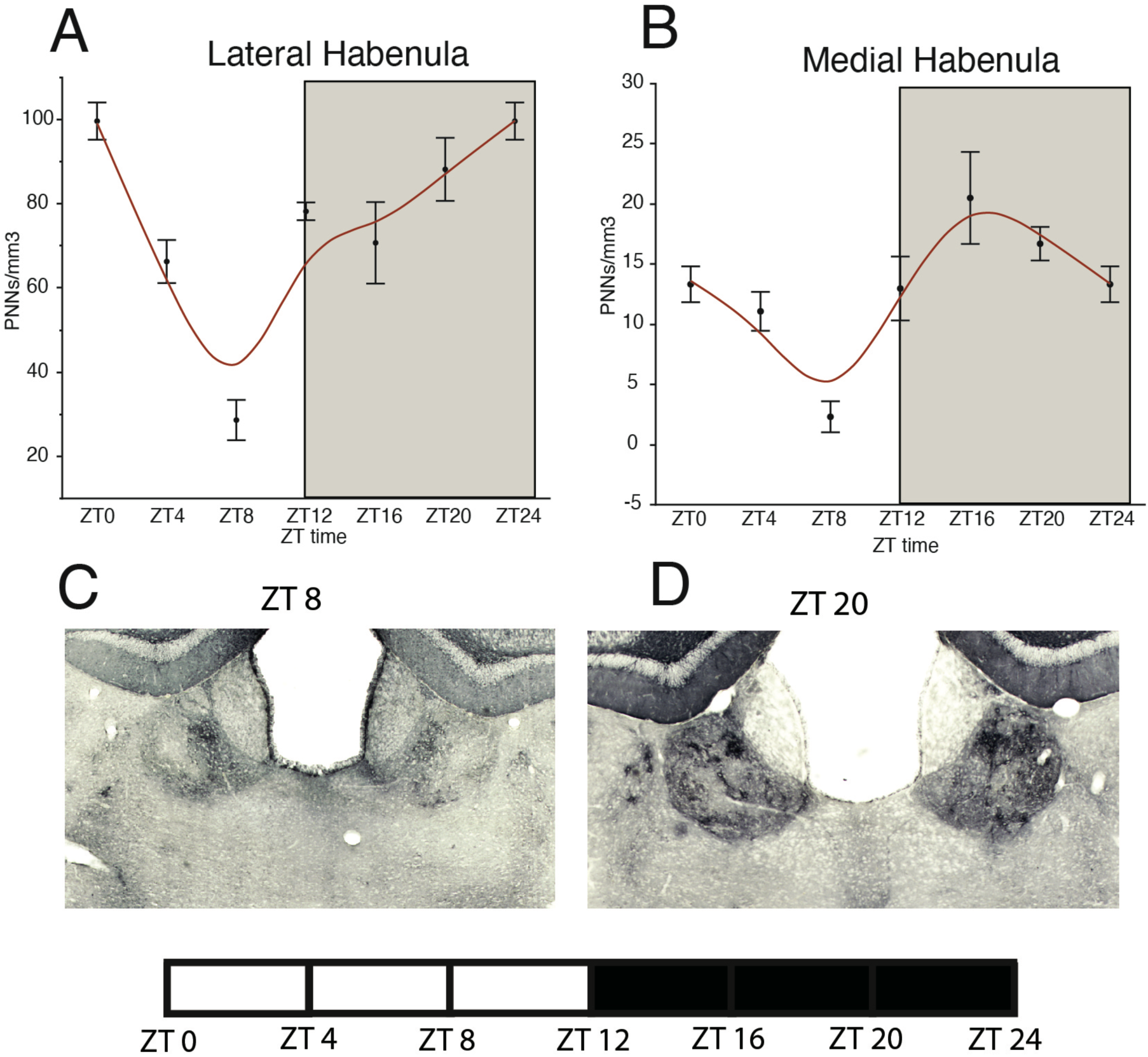
Diurnal Rhythms of Perineuronal Nets in the Mouse Habenula. Analysis of WFA+ PNNs across the 24 hour cycle in male mice housed in a 12:12 light/dark cycle revealed a diurnal rhythm of WFA+ PNNs in the lateral habenula (A) and medial habenula (B), with a peak at ∼ZT 0 and trough at ∼ZT 8 for the lateral habenula, and a peak at ∼ZT 16 and trough at ∼ZT 8 for the medial habenula. Error bars represent standard deviation. Representative low magnification images of WFA labeling in the mouse habenula at ZT 8 (C) and ZT 20 (D).

**Figure 5:**
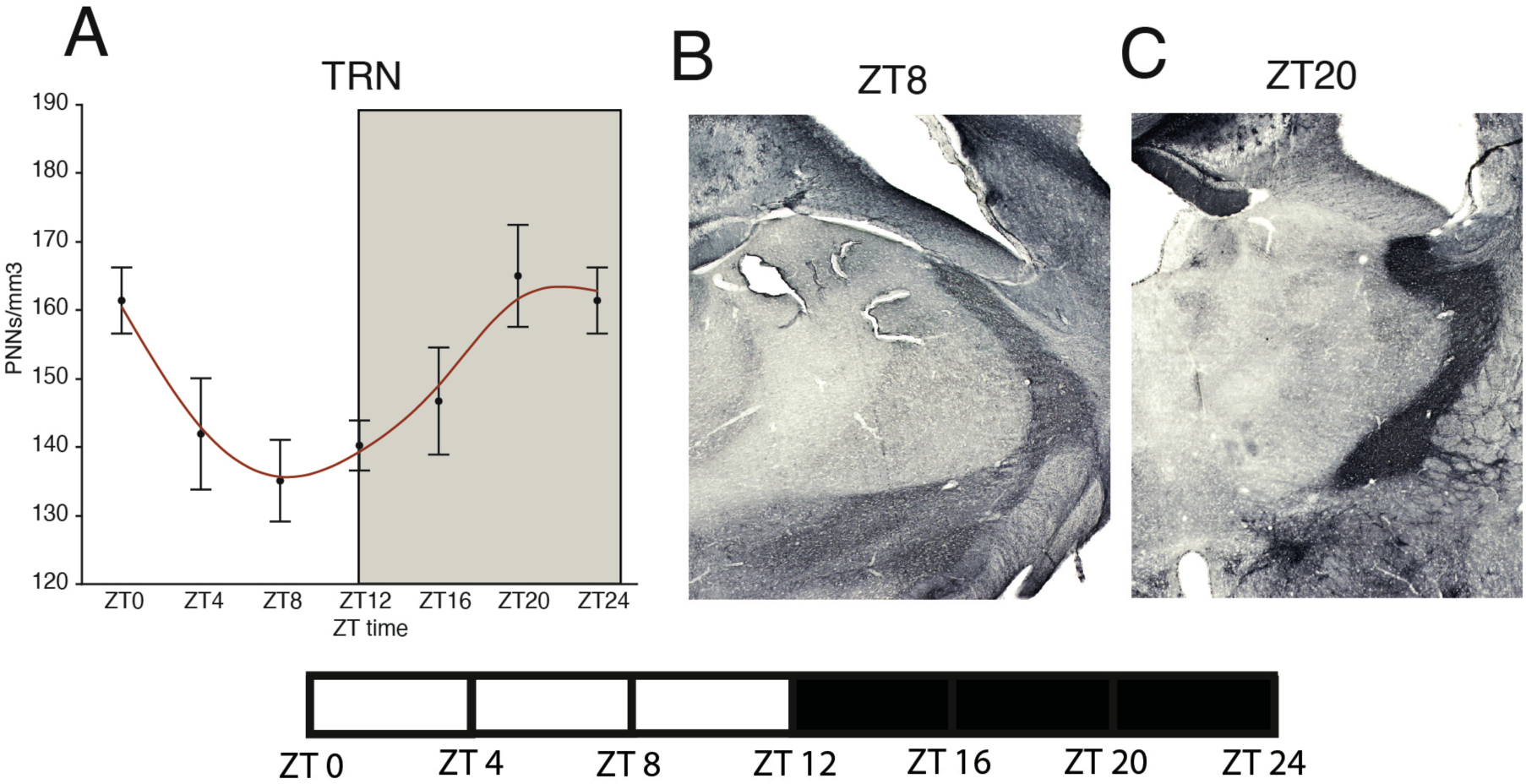
Diurnal Rhythms of Perineuronal Nets in the Mouse Thalamic Reticular Nucleus. Analysis of WFA+ PNNs across the 24 hour cycle in male mice housed in a 12:12 light/dark cycle revealed a diurnal rhythm of WFA+ PNNs in the thalamic reticular nucleus (A) with a peak at ∼ZT 20 and a trough at ∼ZT 8. Error bars represent standard deviation. Representative low magnification images of WFA labeling in the mouse thalamic reticular nucleus at ZT 8 (B) and ZT 20 (C).

**Figure 6:**
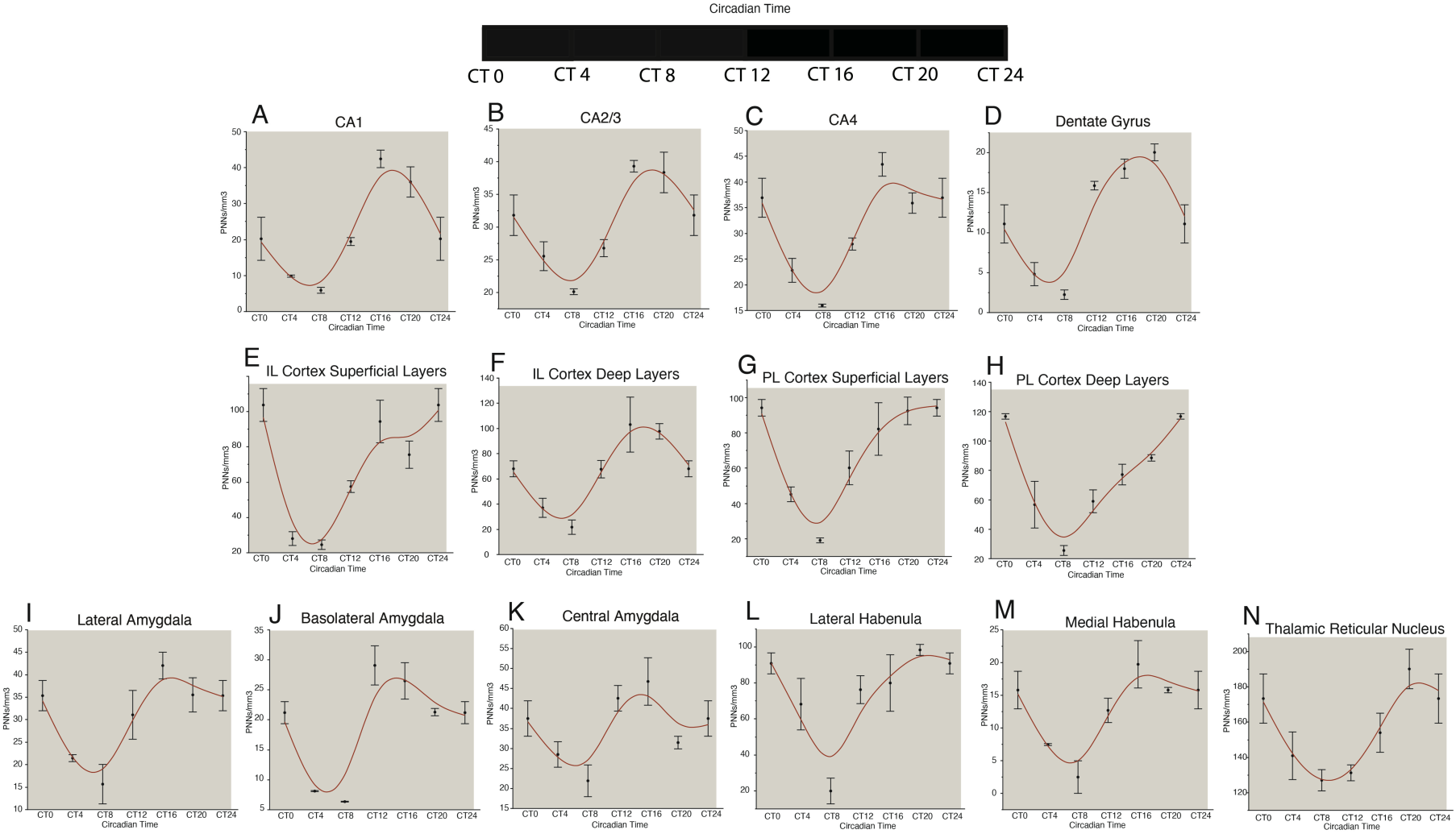
Circadian Rhythms of Perineuronal Nets in the Mouse Brain. Circadian rhythms in the density of WFA+ PNNs were observed in mice housed in constant darkness. In the hippocampus, these rhythms were similar to the diurnal rhythms observed in the CA regions and the DG (A-D), Circadian rhythms in the density of WFA+ PNNs in the mouse prefrontal cortex also paralleled the observed diurnal rhythms in these regions, with peaks at ∼CT 0 and troughs at ∼CT 8 (E-H), with the exception of the deep layers of the IL cortex, which showed a peak at ∼CT 20 and trough at CT 8 (F). Circadian rhythms of WFA+ PNN densities were also observed in the lateral, basal, and central amygdala nuclei in constant darkness, with a peak at ∼CT 16 and a trough at ∼CT 6 (I-K). Circadian rhythms of WFA+ PNN densities in the lateral and medial habenula and thalamic reticular nucleus paralleled diurnal PNN rhythms in these regions (L-N). Error bars represent standard deviations.

### Circadian Rhythms of PNNs in the Mouse Brain

These studies were designed to assess whether diurnal rhythms in mice reflect a true circadian rhythm, and to confirm the existence of PNN density rhythms in a separate strain of mice. We used adult male C57/Bl6 mice housed in a 12:12 LD cycle and then placed into constant darkness for three full 24-hour cycles in order to quantify WFA+ PNN rhythms in free-running circadian conditions. In mice kept in constant darkness, numerical density of WFA+ PNNs displayed circadian rhythms in all regions identical to diurnal rhythms described above, with consistent peaks at approximately CT20 and troughs at approximately CT8 across regions (Fig.6).

### Sleep Deprivation Prevents the Decrease of PNNs During the Day in the Mouse Hippocampus

Sleep deprivation by gentle handling has been previously shown to prevent synaptic modification that occurs in the hippocampus during sleep in rodents (Havekes et al., 2016; Raven et al., 2018). Here, we use the same approach to test the hypothesis that sleep deprivation prevents the decrease of WFA+ PNN densities in the mouse hippocampus. Mice that were sleep deprived for 5 hours (ZT0-ZT5) starting from the beginning of the light cycle had significantly higher numerical density of WFA+ PNNs in the dentate gyrus (p = 0.01) and sectors CA4 (p= 0.01), CA3/2 (p= 0.01) and CA1 (p= 0.001) (Fig. 7B-D). Similar differences in WFA+ PNN densities were observed in the amygdala, habenula, and prefrontal cortex (Fig. 7E-G). In a set of animals that underwent auditory fear conditioning, 5 hours of sleep deprivation significantly enhanced fear memory extinction (Fig. 7A).

**Figure 7:**
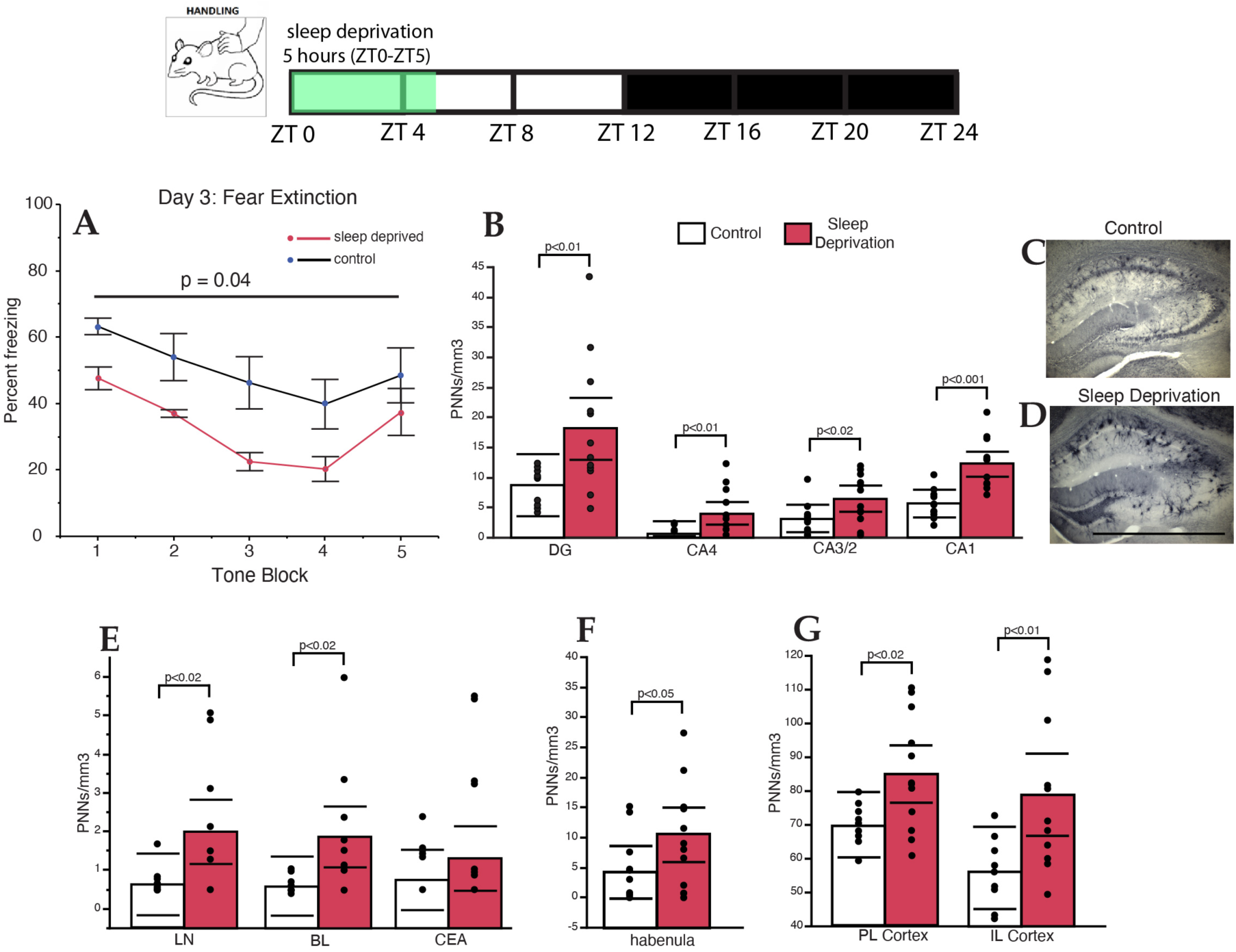
Sleep Deprivation Prevents PNN Decreases. Five hours of sleep deprivation, from lights on (7AM) to 12 PM following auditory fear conditioning, resulted in rapid extinction of fear memory (A), along with significantly higher numerical density of WFA+ PNNs in the hippocampus (B). Representative photomicrographs of the hippocampus labeled with WFA from a control mouse (C) and a sleep deprived mouse (D). Scale bar = 1000 μm. Similar increases in densities of WFA+ PNNs in SD mice were also observed in the amygdala (E) habenula (F) and prefrontal cortex (G). Error bars represent 95% confidence intervals.

**Figure 8:**
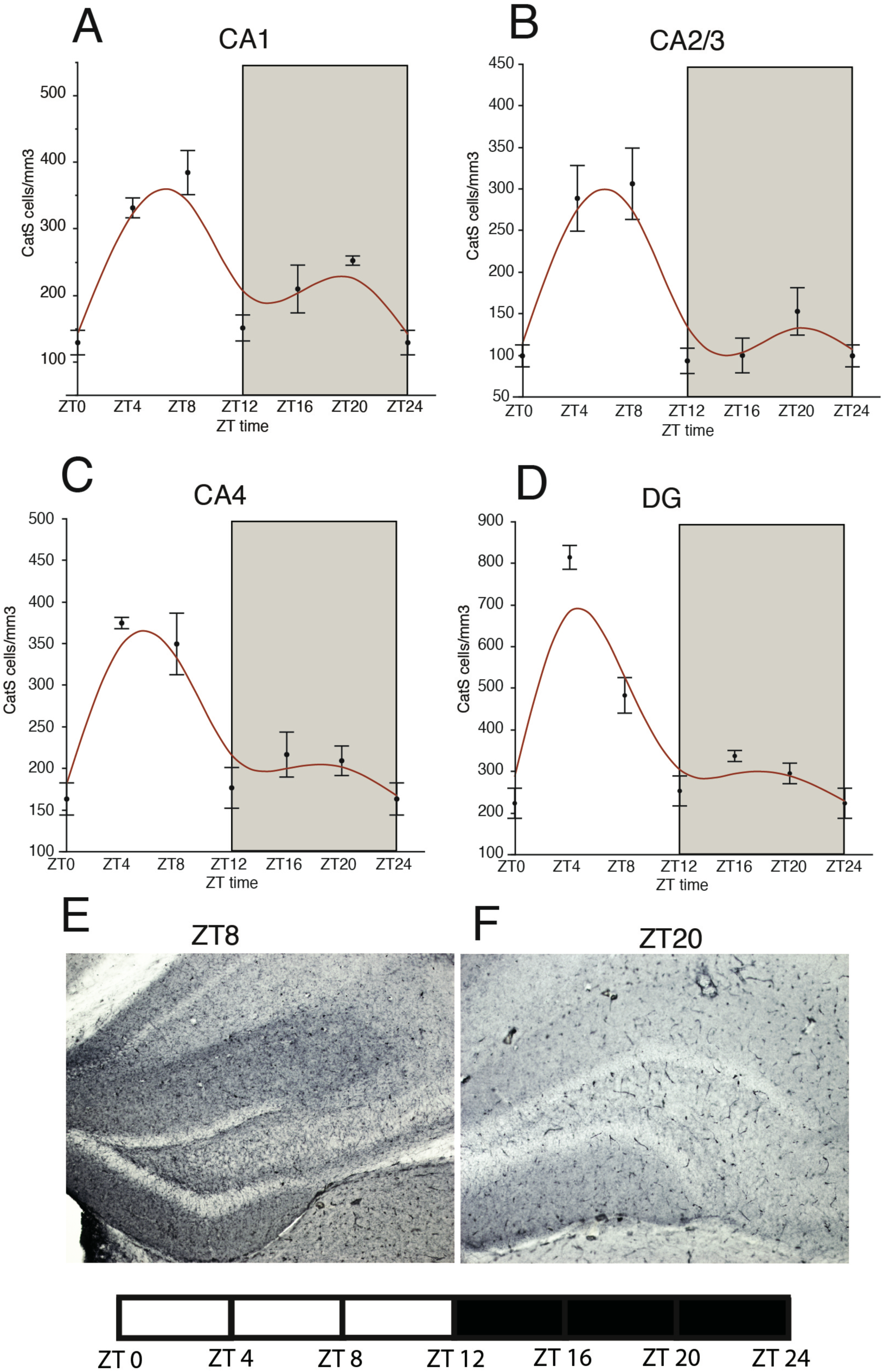
Cathepsin-S Diurnal Rhythms in the Mouse Hippocampus. Diurnal rhythms in densities of cathepsin-S immunoreactive cells were observed in CA1 (A), CA2/3 (B), CA4 (C), and the DG (D) in mice, with expression peaking during the middle of the light cycle, when WFA+ PNN numbers are low in these regions, and decreasing during the dark cycle, when WFA+ PNN densities are high. Error bars represent standard error of the mean. Representative photomicrographs of the hippocampus labeled with cathepsin-S at ZT 8 (E) and ZT 20 (F).

### Association of Cathepsin-S Microglia with Diurnal PNN Rhythms

Cathepsin-S has been reported to be rhythmically expressed in the mouse prefrontal cortex and associated with diurnal rhythms in dendritic spines and electrophysiological properties of prefrontal cortex neurons (Hayashi et al., 2013b). As a first step in testing whether cathepsin-S may contribute to circadian modification of PNN integrity, we tested the hypothesis that its expression in microglia may vary according to a diurnal rhythm, antiphase to WFA+ PNN rhythms. We observed a diurnal rhythms of cathepsin-S-immunoreactive microglia densities in the mouse hippocampus, antiphase to the rhythms of WFA+ PNNs in this region (Fig. 8), with peaks at approximately ZT6 and troughs at ZT0. Similar diurnal cathepsin-S rhythms were observed in the amygdala and prefrontal cortex (Fig.9). No significant effects of average daily wheel-running activity on cathepsin-S immunoreactive cell densities were observed (Table 3). Finally, in order to confirm that cathepsin-S degrades PNNs, we incubated mouse sections in active cathepsin-S (3 hours and 24 hours). Our results show an incubation time-dependent decrease of WFA+ PNN labeling, with a significant 54.5% decrease after 3 hours (p<0.02) and a complete elimination after 24 hours (p<0.0001) (Fig. 10 A-F). Dual immunohistochemistry confirmed that virtually all (88.12-92.95%) of cathepsin-S immunoreactive cells in the mouse hippocampus, infralimbic and prelimbic cortex, amygdala, habenula, and TRN correspond to IBA1 positive microglia (Fig. 10 G-N).

**Figure 9:**
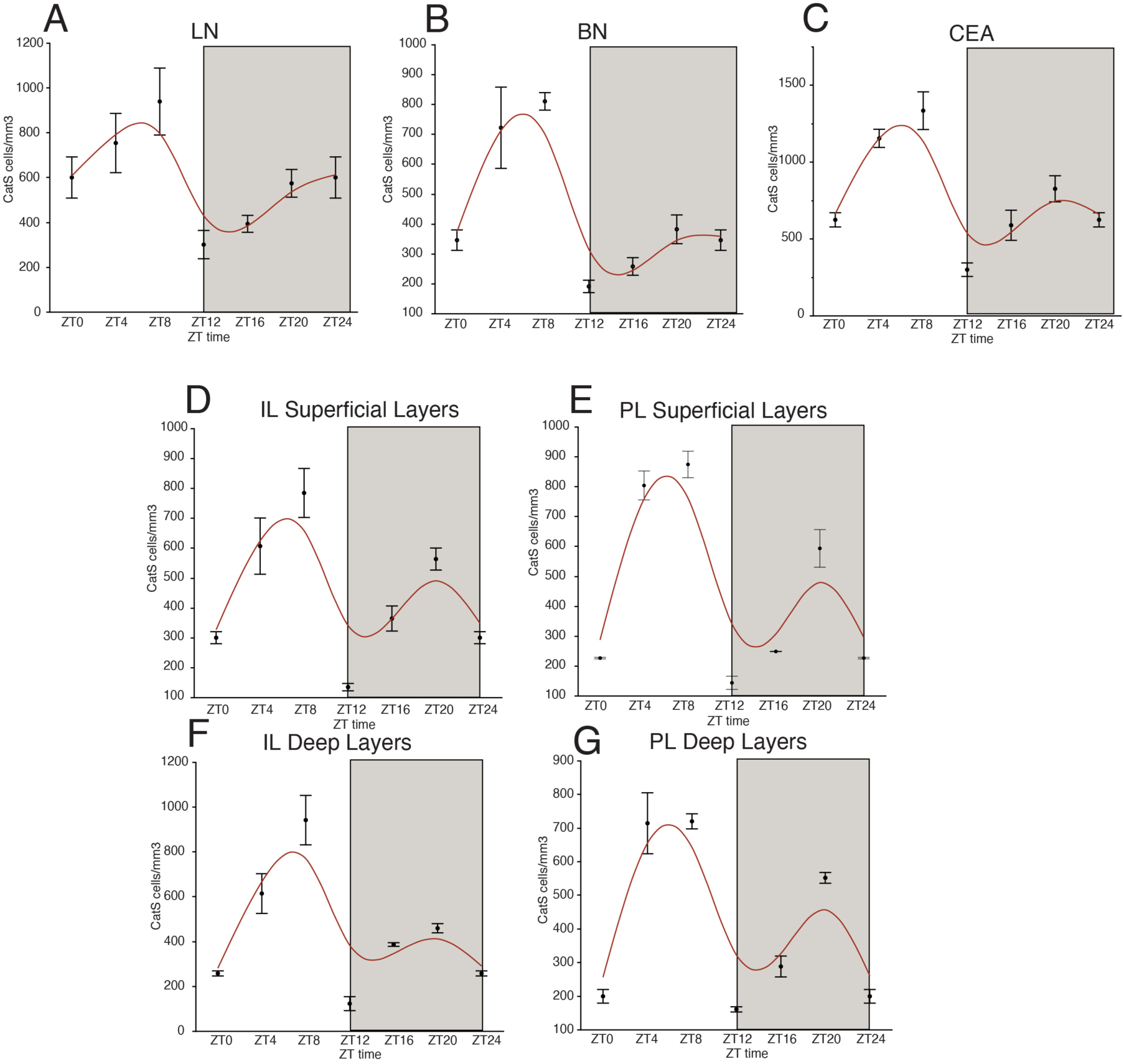
Cathepsin-S Diurnal Rhythms in the Mouse Amygdala and Prefrontal Cortex. Diurnal rhythms in densities of cathepsin-S immunoreactive cells were observed in the lateral amygdala (A), basal amygdala (B) and central amygdala (C), with expression peaking during the middle of the light cycle, when WFA+ PNN numbers are low in these regions, and decreasing during the dark cycle, when WFA+ PNN densities are high. Similar diurnal rhythms were also observed in the infralimbic cortex superficial layers (D), prelimbic cortex superficial layers (E), infralimbic cortex deep layers (F), and prelimbic cortex deep layers (G). Error bars represent standard error of the mean.

**Figure 10:**
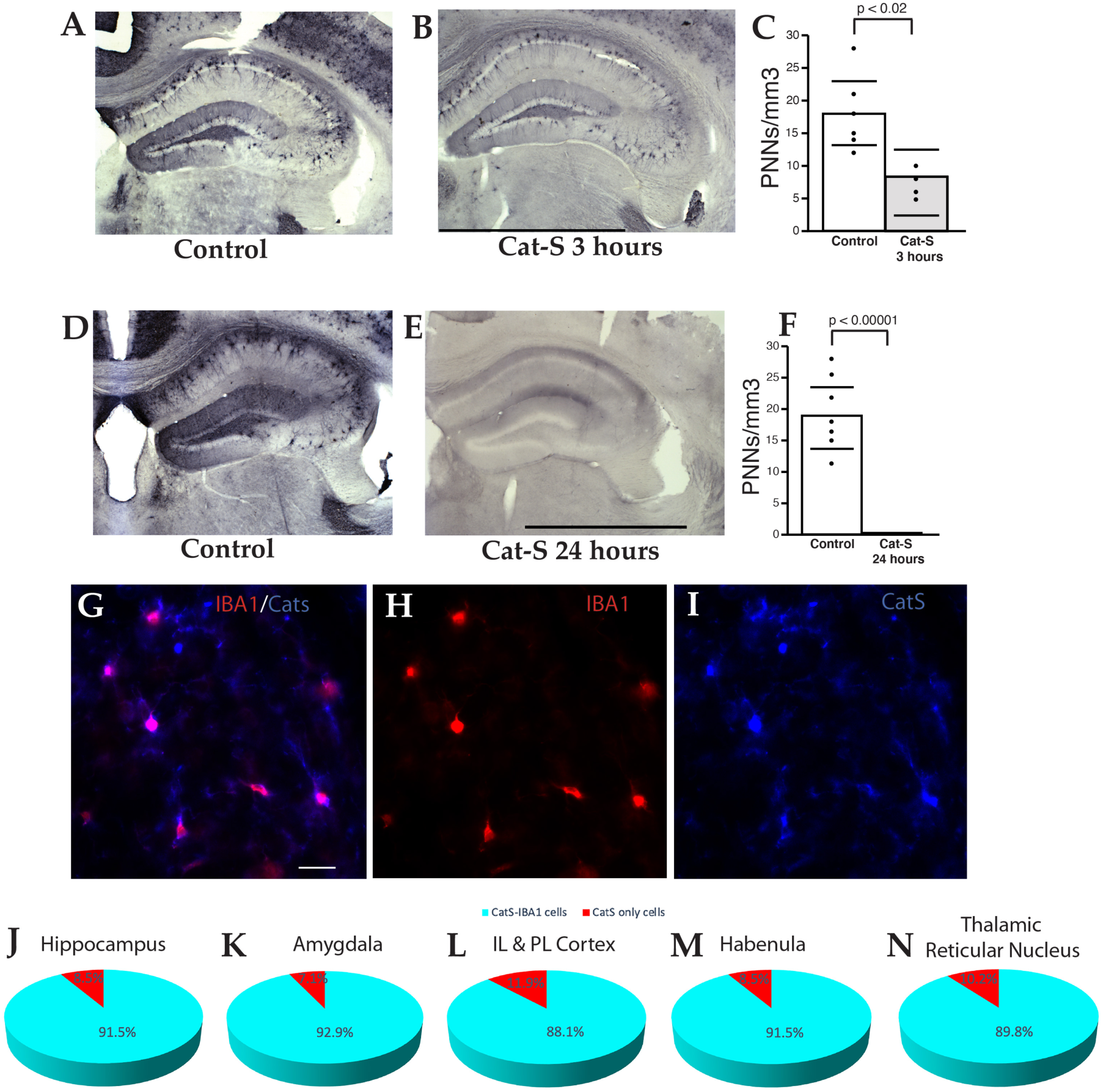
Cathepsin-S is Expressed in Microglia and Eliminates PNN Labeling. Significant reduction in WFA+ PNNs is observed after 3 hours of cathepsin-S incubation (A-C), and a complete absence of PNN labeling after 24 hours (D-F). Error bars represent 95% confidence interval. Scale bars = 1000 μm. Dual fluorescence immunohistochemistry demonstrated that the vast majority of cathepsin-S immunoreactive cells in the mouse hippocampus co-express the microglial marker IBA1 (G-N). Scale bar = 50 μm.

### Diurnal Rhythms of PNNs in the Human Amygdala and TRN

For these studies, we used time of death (TOD) for each subject as a <u>proxy for diurnal rhythms (zeitgeber time; see Discussion)</u> to test the hypothesis that WFA+ PNN numbers vary in a diurnal manner in the human amygdala and TRN. We observed differences in WFA+ PNN numbers in subjects with a TOD during the day in comparison to subjects with a TOD during the night in the human amygdala (Fig. 11 A-D). Quartic regression analysis revealed a diurnal rhythm of WFA+ PNNs Nt in the human amygdala (Fig. 11D), with peaks at noon and midnight, and troughs at 4 AM and 8 PM. In contrast, we observed day/night differences in WFA+ PNN numbers in the human TRN that are opposite to the human amygdala, with higher numbers of PNNs at night and lower numbers during the day (Fig. 11 E-H). Quartic regression plots revealed peaks of WFA+ PNN numbers in the TRN at night during 4 AM and 8 PM, and the lowest numbers at 12 PM and midnight (Fig. 11 H).

**Figure 11:**
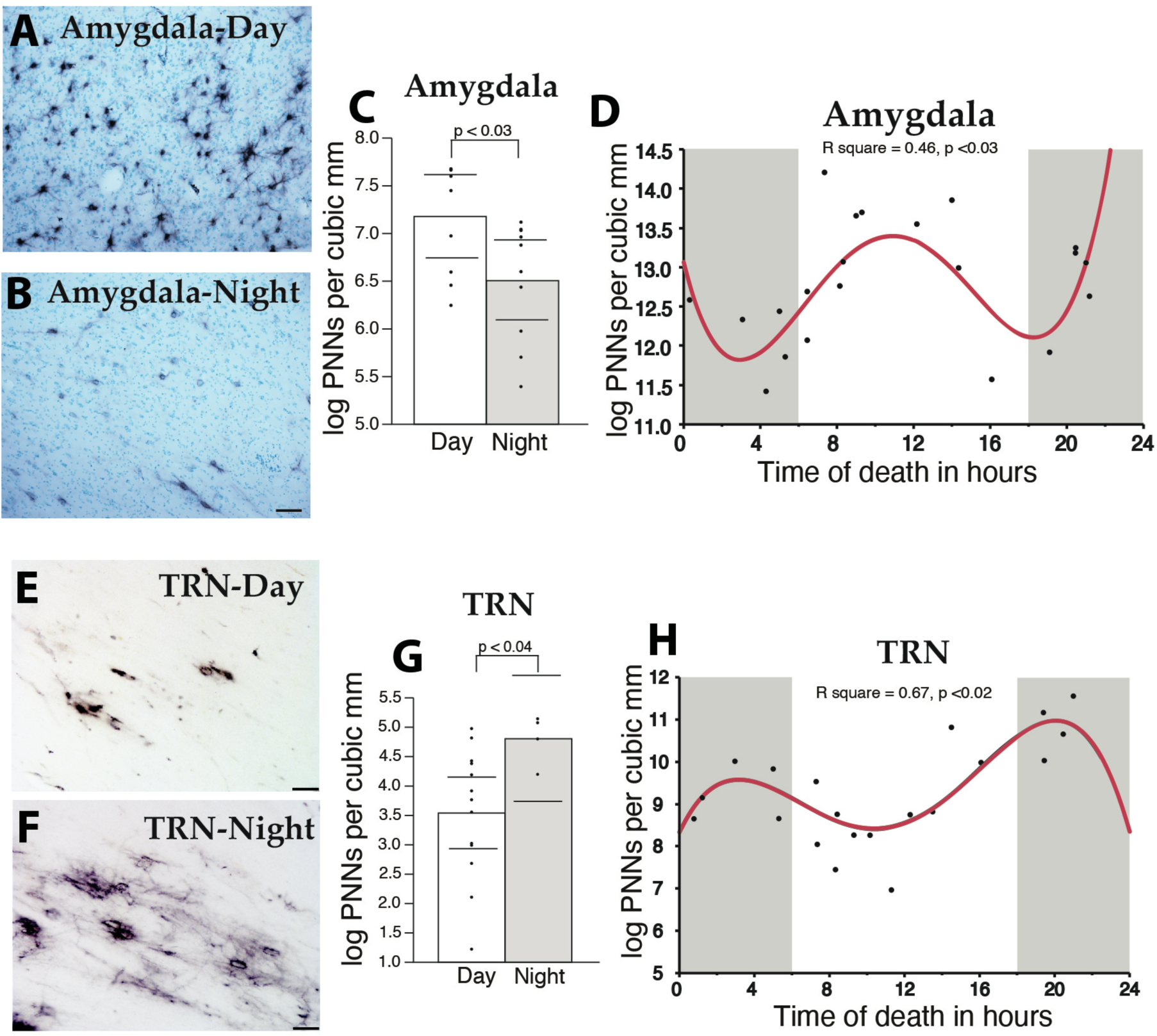
Diurnal Rhythms of Perineuronal Nets in the Human Brain. WFA+ PNN numbers vary with time of death in the human brain. Photomicrograph depicting PNN labeling by WFA lectin in the human amygdala during the day (A) and at night (B). WFA+ PNNs displayed a significant day/night difference in the human amygdala (C), with peaks PNN numbers at noon and midnight, and troughs at 4 AM and 8 PM (D). Photomicrograph depicting PNN labeling in the human thalamic reticular nucleus (TRN) during the day (E) and at night (F). Significant day/night differences were observed in total numbers of WFA+ PNNs in the TRN (G). Quartic regression plots revealed a dual peak rhythm in the TRN that is antiphase to the rhythm observed in the amygdala (H). Error bars represent 95% confidence intervals.

## Discussion

We present, to our knowledge for the first time, evidence that WFA+ PNN vary according to diurnal rhythms in the human brain and to diurnal and circadian rhythms in the rodent brain. Our data adds to a growing number of studies demonstrating that PNNs are dynamic structures, responding to the environment and potentially contributing to memory consolidation during sleep (Balmer et al., 2009; Brown et al., 2009; Banerjee et al., 2017; Dingess et al., 2018; Slaker et al., 2018), We show that numbers of WFA+ PNN follow diurnal rhythms in several brain regions in mouse and in human. Importantly, we show that WFA+PNN rhythmicity occurs in mice kept in constant darkness, supporting the claim that these changes reflect circadian rhythms rather than a response to light-dark cycles (Fig.6). Our results also provide evidence for a role of the microglia-derived matrix protease cathepsin-S, known to contribute to synaptic plasticity (Hayashi et al., 2013b). We show a diurnal rhythm of cathepsin-S expression in microglia, opposite to the observed PNN rhythms, and demonstrate that cathepsin-S eliminates WFA+ PNN labeling. Taken together, these results support the hypothesis that cathepsin-S may represent one of the endogenous proteases contributing to WFA+ PNN rhythms. PNN rhythms in the mouse hippocampus coincide with reported rhythms in LTP, suggesting that WFA+ PNNs decrease during sleep, when lower levels of LTP were reported to occur (Chaudhury et al., 2005), and increase during wakefulness when higher levels of LTP occur as animals encode new memories (Hou et al., 2013) (Fig. 6). We suggest that diurnal rhythms of WFA+ PNNs in the regions examined may have broad implications for emotional memory processing and psychiatric disorders.

### Technical Considerations

#### Interpretation of WFA+ PNN rhythms

PNNs are highly complex structures formed by several glycoproteins and proteoglycans, link proteins and hyaluronan (Maeda, 2010; Miyata et al., 2012). WFA detects a specific sulfation motif on N-acetylgalactosamine at the terminal ends of CS chains (Caterson et al., 1990; Miyata et al., 2012). CS chains can be modified by addition of sulfation groups at the 2, 4 or 6 positions along the chains, allowing for highly complex and dynamic modification (Caterson et al., 1990; Maeda, 2010; Miyata et al., 2012; Pantazopoulos et al., 2015). Furthermore, CS chains can be cleaved at varying points along the chain by several matrix proteases (Muir et al., 2002; Porter et al., 2005; Pantazopoulos et al., 2015). Together, these considerations suggest that it is unlikely that the complex PNN structure may be entirely degraded and rebuilt on a 24-hour cycle. We propose that the diurnal and circadian WFA+ PNN rhythms we observed may reflect modifications of the biochemical characteristics of these structures, perhaps impacting the CS chain sulfation pattern detected by WFA. It is important to emphasize that growing and compelling evidence shows that PNN and ECM functions are dictated by dynamic posttranslational modifications of their components mediated by matrix proteases (Pantazopoulos and Berretta, 2016; Lasek et al., 2018; Wen et al., 2018). Notably, these modifications determine whether effects of ECM components on synaptic plasticity are inhibitory or permissive (Miyata et al., 2012; Foscarin et al., 2017; Yang et al., 2017). Our data showing cathepsin-S rhythms antiphase to PNN rhythms, and the ability of cathepsin-S to eliminate WFA+ PNN labeling, support this interpretation and represent the first step in examining this process. However, a significant limitation is that our current data show associations but do not demonstrate mechanistic effects of cathepsin-S expression rhythms on PNN rhythms. Our data showing diurnal rhythms of cathepsin-S expression represents the first step in testing a broad range of proteases from several cell types. Circadian regulation of PNNs is likely to consist of a complex molecular signaling system involving multiple proteases and ECM molecules, encompassing several cell types. Future studies focused on circadian expression of specific PNN components, matrix proteases and sulfotransferases will provide insight into the mechanisms underlying circadian PNN modification and direct effects on memory processing.

#### Time of death (TOD) in human postmortem subjects as a proxy for diurnal rhythms (zeitgeber time)

Human postmortem studies have successfully used TOD as a proxy for diurnal rhythms (approximate zeitgeber time), to study diurnal rhythms of gene and protein expression in human brain. An obvious limitation is that TOD represents a single measure per subject at a specific time point, rather than repeated measures across time. However, several human studies demonstrated predicted rhythmic expression of clock genes in several brain regions, and of SST in the amygdala (Li et al., 2013; Bunney et al., 2015; Chen et al., 2016; Pantazopoulos et al., 2016). Importantly, rhythmic patterns, such as peak phase relationships between clock molecules, demonstrated in human were consistent with those reported in rodents, including staggered phase relationship between Per1, Per2, and Per3 genes (Lamont et al., 2005; Ramanathan et al., 2010; Albrecht et al., 2013; Li et al., 2013). Molecular rhythms reported in the human cortex have been independently replicated, providing further support for the validity of this approach (Li et al., 2013; Chen et al., 2016). The WFA+ PNN rhythms observed in the amygdala nocturnal mice (Figs. 1 & 6) are antiphase to the PNN rhythms observed in diurnal human subjects in the same region (Fig. 11), providing further support for the approach of using TOD to analyze rhythmic relationships in human postmortem samples.

### Implications for Synaptic Plasticity and Memory Consolidation

Several hypotheses have been put forth to link wake/sleep cycles to synaptic mechanisms underlying memory consolidation. For instance, studies from Tononi and Cirelli support the synaptic homeostasis hypothesis of sleep (Tononi and Cirelli, 2006, 2014). Briefly, neurons form and strengthen many new synapses during wakefulness, as organisms interact with their environment and encode new memories. During sleep, when the active encoding process is offline, synapses are downscaled, in order to enhance the signal to noise ratio, thus improving memory function (Tononi and Cirelli, 2006, 2014). Consistent with this hypothesis, decreases of dendritic spines and synapses during sleep have been reported in sensory and motor cortical regions (Maret et al., 2011; de Vivo et al., 2017). An alternative theory, suggested by Rasch and Born, postulates that memories are reorganized during slow wave sleep in a process called systemic consolidation (Rasch and Born, 2013). During systemic consolidation, memory representations are reactivated, and transferred from short-term storage sites such as the hippocampus, into long-term storage in neocortical areas where they are integrated into existing schemas (Rasch and Born, 2013). Memories are then strengthened in these long-term storage areas during REM sleep, in a process called synaptic consolidation, while the short-term storage memories are removed via synaptic pruning (Rasch and Born, 2013).

We speculate that diurnal molecular modifications of PNNs may contribute to memory formation and consolidation mechanisms during the wake/sleep cycle, favoring activity-driven synaptogenesis and synaptic refinement, respectively. For instance, our results on the effects of 5 hour sleep deprivation on WFA+ PNNs in the mouse hippocampus are consistent with reports that 5 hours of sleep deprivation prevents changes in dendritic spine densities in the hippocampus occurring during sleep (Havekes et al., 2016; Raven et al., 2018; Spano et al., 2019; Gisabella et al., 2020). PNN rhythms observed in our study may reflect ongoing systemic and synaptic consolidation during sleep proposed by Rasch and Born (Rasch and Born, 2013). For instance, WFA+PNNs changes in mice the hippocampus are more active during wakefulness, as suggested by enhanced LTP in this region during the night (Chaudhury et al., 2005). Regional differences in PNN rhythms may also reflect phase differences in molecular clock rhythms of these regions. Region specific rhythms in the clock protein Per2 have been described previously in rodents and humans (Lamont et al., 2005; Li et al., 2013; Harbour et al., 2014; Chen et al., 2016).

Recent evidence shows that cathepsin-S deletion in knockout mice contributes to failure to downscale synapses during sleep (Hayashi et al., 2013b). In these mice, reduced EEG delta wave power, failure to reduce amplitude and frequency of action potentials and to reduce dendritic spines during sleep supports a role for cathepsin-S in downscaling synaptic strength during sleep (Hayashi et al., 2013b). Our results show that rhythms of cathepsin-S expression are antiphase with respect to WFA+ PNNs rhythms, i.e. high cathepsin-S expression is associated with low WFA+ PNN numbers, and that cathepsin-S reduces WFA+ PNN labeling. Together, these findings suggest that increased cathepsin-S during sleep may represent one of several molecules that contribute to modifying PNN composition. In turn, such modifications may contribute to synaptic downscaling and remodeling during memory consolidation (Fig. 12). This hypothesis is supported by evidence for the involvement of the microglial circadian molecular clock in the regulation of microglial morphology, immune response, and synaptic regulation (Hayashi et al., 2013b; Hayashi et al., 2013a; Fonken et al., 2015). Our results suggest an additional circadian role for microglia in synaptic regulation, through PNN modification potentially modulating memory consolidation processes.

**Figure 12:**
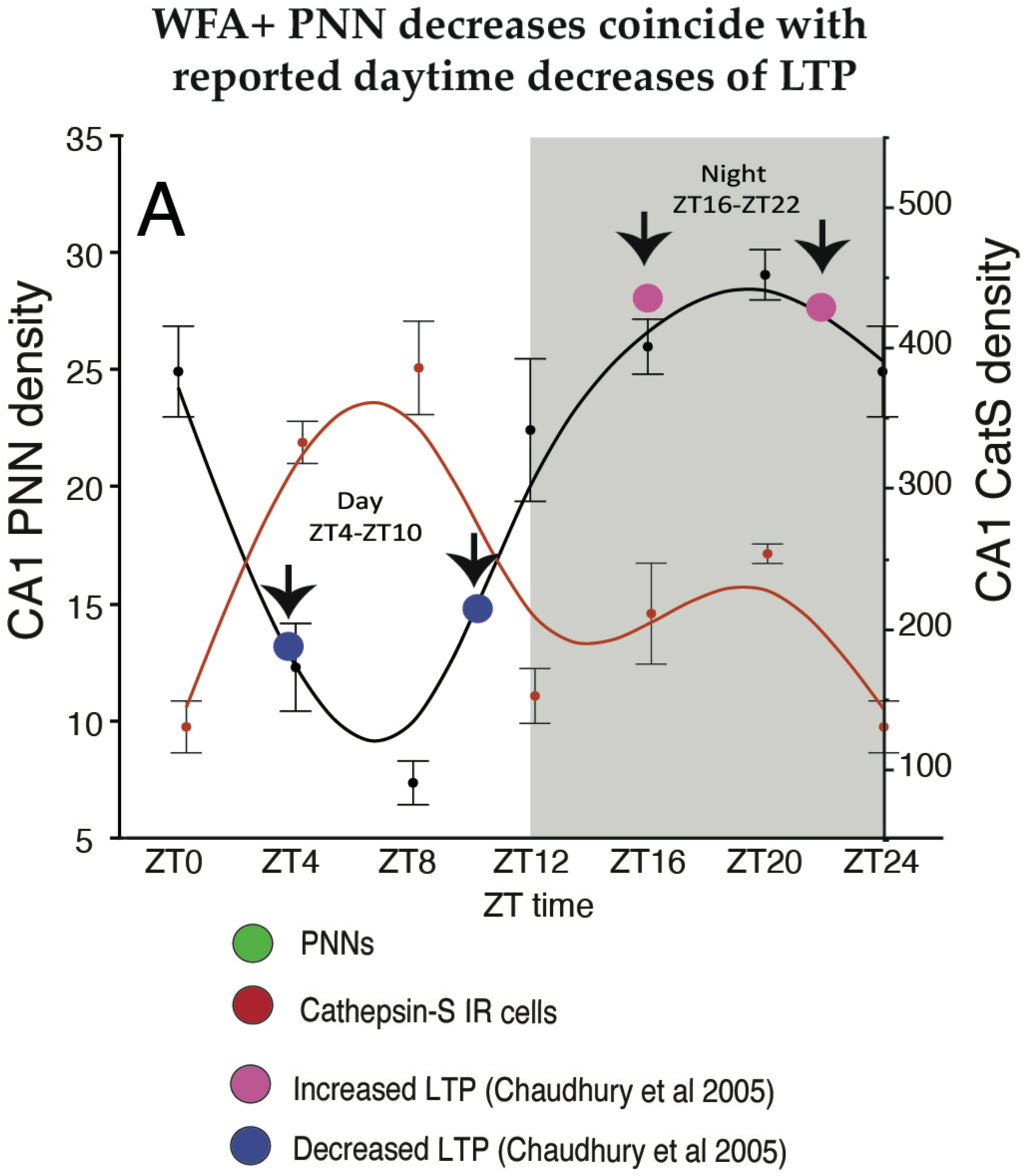
Microglial Expression of Cathepsin-S May Modify PNNs to Allow for Memory Consolidation During Sleep. In the mouse hippocampal sector CA1, diurnal rhythms in the numerical density of WFA+ PNNs decreases during the day as mice sleep, reaching the lowest density in WFA+ PNN numbers between ZT 4-ZT 10 (green curved line). This coincides with the peak expression of cathepsin-S (red curved line) and the reported daytime decrease in LTP (blue circles, from Chaudhury et al 2005). In comparison, the numerical density of WFA+ PNNs peaks during the dark at ∼ZT 20 during the active period for nocturnal mice, coinciding with the low point of cathepsin-S immunoreactivity in this region as well as the reported increase in LTP at night in mice (pink circles, from Chaudhury et al 2005). These results suggest that cathepsin-S modifies PNN composition, coinciding with decreased TLP during sleep, to allow for memory consolidation, and PNN composition is restored during the active wake periods to allow for optimal encoding of novel information.

Circadian rhythms in PNN composition may also be regulated by proteases and CSPG production from several cell types including astrocytes and neurons, which produce many of the core PNN components as well as endogenous proteases known to modify PNNs (Pantazopoulos and Berretta, 2016; Miyata and Kitagawa, 2017; Bozzelli et al., 2018). Furthermore, although our evidence suggests that circadian rhythms in PNN composition may contribute to synaptic regulation during sleep, we do not demonstrate a mechanistic effect on synaptic regulation or memory consolidation. PNN circadian rhythms may be involved in other processes such as resolution of oxidative stress during sleep. Several studies suggest that sleep deprivation contributes to increased oxidative stress in in the brain (Silva et al., 2004; Ramanathan and Siegel, 2011; Alzoubi et al., 2012; Harkness et al., 2019), and PNNs are critically involved in protecting fast-firing neurons from oxidative stress (Cabungcal et al., 2013). Thus, rhythms in PNN composition may reflect periods of reduced neuronal activity and resolution of oxidative stress during sleep. A recent study reporting increased oxidative stress in parvalbumin neurons together with increased WFA labeling of PNNs following sleep deprivation supports this hypothesis (Harkness et al., 2019).

### Implications for Psychiatric Disorders

In the present study, we focused on brain regions involved in emotional memory processing and implicated in psychiatric disorders (Vyas et al., 2002; Sartorius et al., 2010; Li et al., 2011; Mahan and Ressler, 2012; Mauney et al., 2013; Meyer et al., 2014; Pantazopoulos et al., 2017; Wells et al., 2017). Diurnal rhythms of PNNs in human subjects have broad implications for psychiatric disorders. PNN deficits have been reported by several groups in the amygdala, entorhinal cortex, hippocampus, prefrontal cortex, and TRN in schizophrenia and bipolar disorder (Pantazopoulos et al., 2010a; Mauney et al., 2013; Pantazopoulos et al., 2014; Pantazopoulos et al., 2015; Enwright et al., 2016; Steullet et al., 2017). Disruption of PNNs in these disorders may alter rhythms of synaptic plasticity and in turn contribute to shared synaptic deficits (Penzes et al., 2011; Glausier and Lewis, 2013; Shelton et al., 2015; MacDonald et al., 2017). Such deficits may arise from disrupted memory consolidation processes allowing for decreased synaptic formation and/or increased synaptic pruning in brain regions involved in emotional memory processing.

Abnormalities in sleep and circadian rhythms have also been consistently reported in these disorders (McClung, 2013; Manoach et al., 2016; Pantazopoulos et al., 2017; Seney et al., 2019). Decreased sleep spindles, generated by the TRN, and memory consolidation deficits are emerging as consistent characteristics of schizophrenia (Ferrarelli et al., 2007; Manoach et al., 2010; Manoach et al., 2014). Decreased sleep spindles have been reported in several independent studies, including in unmedicated patients with schizophrenia, and in first-degree relatives, suggesting that this represents a core genetic component of the disease rather than medication effects or consequence of disease progression (Ferrarelli et al., 2007; Manoach et al., 2010; Manoach et al., 2014). Disruption of WFA+ PNN rhythms in subjects with schizophrenia may contribute to sleep spindle and memory consolidation deficits in several ways. WFA+ PNNs regulate firing rates of neurons expressing parvalbumin (PVB), including those in the TRN that generate sleep spindles (Csillik et al., 2005; Katsuki et al., 2017). Furthermore, decreases of PVB neurons were detected in the TRN of subjects with schizophrenia (Steullet et al., 2017). PNNs protect PVB neurons from oxidative stress (Cabungcal et al., 2013), thus disruption of PNN rhythms may leave PVB neurons more susceptible to accumulation of oxidative damage during sleep, resulting in loss of PVB neurons in subjects with schizophrenia (Steullet et al., 2017). PVB deficits in TRN function have been proposed by several groups to contribute to memory consolidation deficits in schizophrenia (Manoach et al., 2016; Ferrarelli and Tononi, 2017). Disrupted PNN rhythms in the TRN may contribute to a decreased ability of this region to generate sleep spindles, and in turn memory consolidation deficits. In addition, disrupted PNN rhythm composition by cathepsin-S in expression from microglia in subjects with schizophrenia may contribute to memory consolidation deficits through disruption of local synaptic downscaling and reorganization proposed to occur during sleep (Tononi and Cirelli, 2006; Rasch and Born, 2013; Tononi and Cirelli, 2014). Cathepsin-S knockout mice, in which diurnal rhythms of dendritic spine density were reported (Hayashi et al., 2013b), also display deficits in social interaction and novel object recognition (Takayama et al., 2017), supporting the hypothesis that cathepsin-S rhythms regulate key roles of PNNs in memory processing and social behaviors that are disrupted in subjects with schizophrenia.

Our findings may also be relevant to the pathophysiology of PTSD. PNNs are strongly involved in fear memory processing, which is enhanced in this disorder (for review see/ (Parsons and Ressler, 2013) (Gogolla et al., 2009; Banerjee et al., 2017). Sleep deprivation has been proposed as an early therapeutic approach for PTSD following a traumatic experience (Kuriyama et al., 2010; Cohen et al., 2012; Cohen et al., 2017). Disruption of molecular processes involved in PNN rhythms may represent one of the potential mechanisms through which sleep deprivation may impact memory consolidation as a possible therapeutic approach for alleviating the strength of fear memories contributing to PTSD.

In summary, we provide evidence for diurnal and circadian rhythms of WFA+ PNN numbers in the human and rodent brain, suggesting that their composition is modified on a daily basis. Rhythms in PNN composition may be mediated in part by cathepsin-S expression originating from microglia. These rhythms may contribute to decreased long-term plasticity reported during sleep in the hippocampus, suggesting a key process through which multiple cell types including microglia modify PNNs to allow for to memory consolidation.

